# Uncovering cell-free protein expression dynamics by a promoter library with diverse strengths

**DOI:** 10.1101/214593

**Authors:** Sabrina Galiñanes Reyes, Yutetsu Kuruma, Soichiro Tsuda

## Abstract

Cell-free protein expression systems have been widely used for synthetic biology and metabolic engineering applications in recent years. Yet little is known about protein expression in the cell-free systems. Here we take a systems approach to uncover underlying dynamics of cell-free protein expression. We construct a set of T7 promoter variants to express proteins at different transcription rates in a reconstituted and *E. coli* extract-based cell-free systems. We find that the maximum expression level and the rate of protein synthesis as responses to the transcription rate change are different in the two cell-free systems, suggesting they are driven by different expression dynamics. We confirm this by constructing a simple mathematical model for each cell-free system, which well reproduce the experimental results and also identify different limiting factors for better protein expression in the two cell-free systems. In particular, they revealed there is a negative feedback effect in the mRNA-protein translation by the PURE system and also identified different limiting factors for better protein expression in the two cell-free systems.

---

Cell-free protein expression systems (cell-free systems) have been increasingly popular for synthetic biology and metabolic engineering applications in recent years^1–3^. They have a wide range of applications from "biomolecular breadboard" to rapidly characterize gene constructs (typically linear DNA constructs)^4^ to express proteins in a micro compartment for *in vitro* evolution^5,6^. There are generally two types of cell-free systems: Extract-based cell-free systems source enzymes necessary for protein expression from crude cell extracts. *Escherichia coli* (*E. coli*)-based system is the most well established system due to the high yield protein expression, but still actively improved for simpler preparation methods^7–11^ and better yield^12,13^. Protein synthesis Using Recombinant Elements (PURE) system is another type of cell-free system^14,15^. In contrast to the extract-based cell-free systems, purified components were reconstituted for protein synthesis. As the concentrations of all the components in the PURE system are known, it is suitable for systematic studies of cell-free protein expression, such as optimization of component concentrations for better protein yield^16^.

Both cell-free systems have been studied extensively over the years, yet little is known about the dynamics of cell-free protein expression. Recent comptuational study pointed out that there are more than 240 components and nearly 1000 reactions are involved in the protein translatin of PURE system^17^. The complex cell-free expression dynamics can be an issue especially when multiple proteins are expressed, such as cell-free reconstitution of Sec translocon^18^ and ATPsynthase^15^, protein complexes as the stoichiometric balance of the synthesized proteins needs to be adjusted for a protein complex to be functional. Although they are usually adjusted by titlating the amount of each DNA input, protein expression levels do not linearly correlate with the amount of DNA^19,20^, especially when the strong T7 promoter is employed for transcription^21^. Because of this nonlinear nature of cell-free protein expression, finding an optimal balance for multiple proteins can be a daunting challenge.

Here we take a systems approach to better understand the dynamics of protein expression in cell-free systems. In particular, we consider a cell-free system as a grey box model (i.e. black box with some prior knowledge)^22^ and study the input-output relationship to infer the internal dynamics of cell-free protein expression. As a prior knowledge, a course-grained model of protein exression in a cell-free system can be described as follows^23–25^:

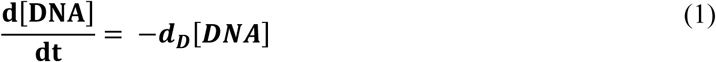

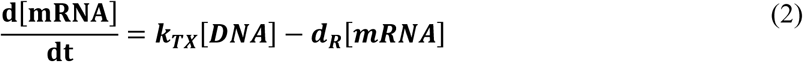

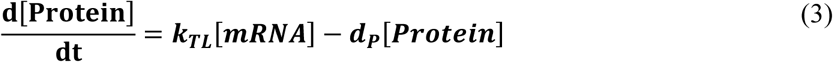

where [DNA], [mRNA] and [Protein] are the concentrations of DNA, mRNA, and protein in the cell-free reaction. *k*_*TX*_ and *k*_*TL*_ are reaction constants for transcription and translation, respectively. *d*_*D*_, *d*_*R*_, and *d*_*P*_ are degradation constants for DNA, mRNA, and protein, respectively.

To estimate the system parameters, system inputs (typically, the concentration of template DNA) were varied and changes in protein expression levels were measured^24^. However, the tunability of system parameters in cell-free systems is not restricted to inputs. It is also possible to modify other parameters by adding/removing system components. For example, addition of GamS (RecBCD nuclease inhibitor) to a crude extract-based cell-free system prevents DNA degradation^4^, which corresponds to decreasing *d*_*D*_ in the equation (1) above.

In this study, we aimed at varying the transcription rate *k*_*TX*_ and measuring changes in protein expression levels as system response. To do this, we constructed variants of the T7 promoter, each of which has a single base-pair (bp) substitutions to the consensus T7 promoter sequence (Figure 1A). It is known that alteration to the consensus sequence affects the binding affinity of T7 RNA polymerase (RNAP) to the promoter and thus changes the protein expression level^26–28^. We evalulated protein expression using the T7 promoter variants in two types of cell-free systems: *E. coli* extract-based system and PURE system. Protein expression dynamics of these two types of cell-free systems are considered to be different owing to lack of supplementary components in the PURE system^14,29,30^. To our best knowledge, however, systematic studies comparing the protein expressions of both cell-free systems have yet to be done. We hypothesized that the two cell-free systems would give different protein expression patterns in response to the varied transcription rates *k*_*TX*_, which would bring new insights to infer the internal dynamics of cell-free protein expression.

**Figure 1.**
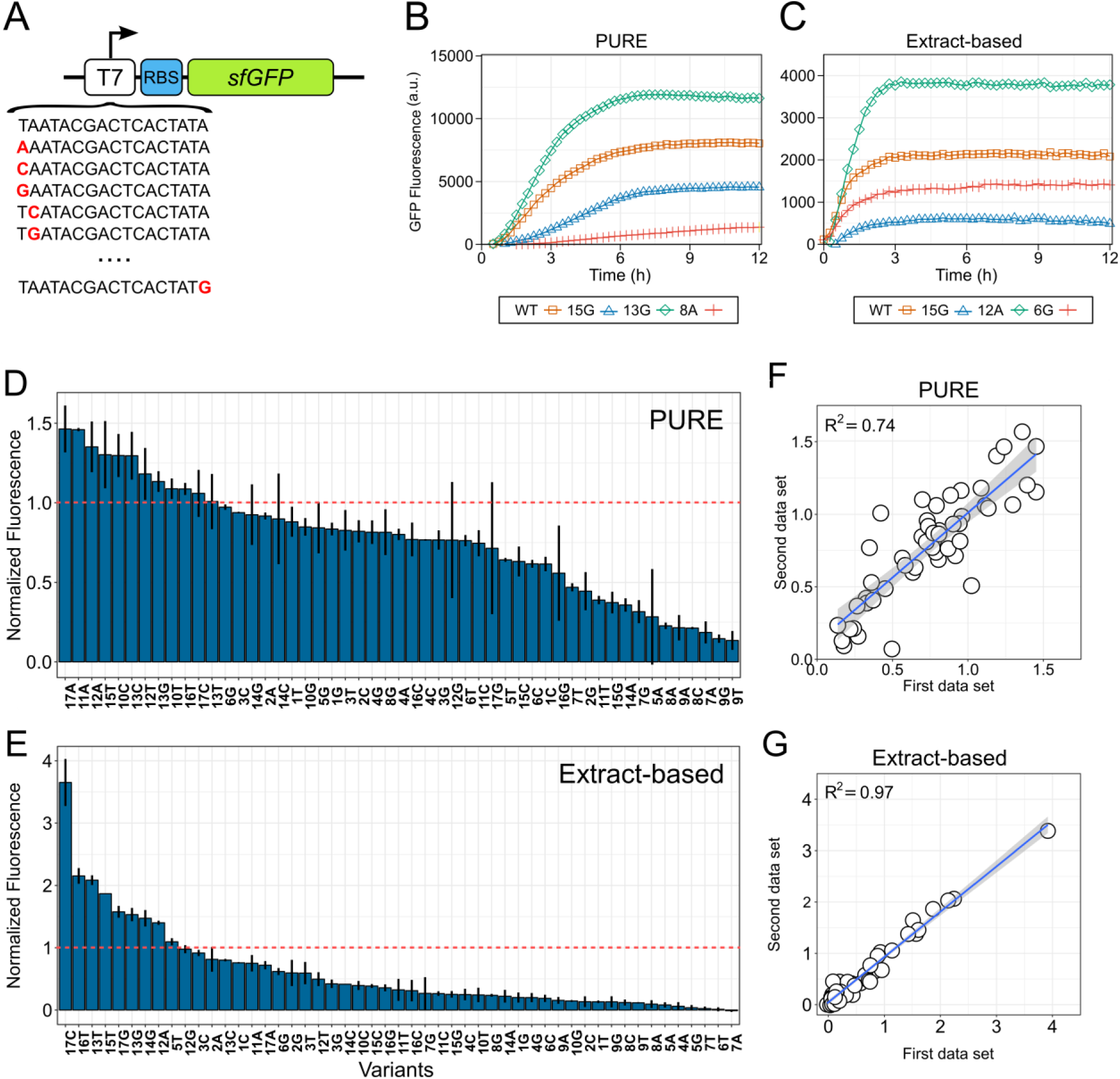
Protein expression in the PURE and extract-based cell-free system using T7 promoter variants. (**A**) A schematic of linear DNA templates (core sequence) used for protein expression. The sequence contained the consensus T7 promoter (or its variants), ribosome binding site (RBS) and the sfGFP gene. Single base-pair substitutions in the T7 promoter variants were highlighted in red. (**B**, **C**) Fluorescence measurements of sfGFP expression with the consensus promoter sequence (denoted as WT) and other three variants in the PURE and extract-based systems, respectively. (**D**, **E**) Relative GFP fluorescence with 51 T7 promoter variants, normalized to that with the consensus sequence (indicated as a red line). Error bars represent the standard deviation. (**F**, **G**) Scatter plots of duplicated experimental data for the PURE and extract-based systems, respectively. The coefficient of determination R^2^ was shown in the upper left part of the plot. The blue line indicates the regression line and the shaded area the 95% confidence interval.

## Results and Discussions

### Characterization of T7 promoter variants in cell-free systems

We systematically altered the recognition site of consensus T7 sequence (TAATACGACTCACTATA) and constructed a library of T7 promoter variants with all possible single base-pair substitutions (Fig.1A). Each variant was ligated with a common ribosome binding site (RBS) and superfolder green fluorescent protein (sfGFP). We hereafter refer to this sequence as "core sequence". Using a total of 51 promoter variants, we investigated the expression level of sfGFP in two types of cell-free systems: *E.coli* extract-based cell-free system (Expressway™ Cell-Free *E. coli* Expression System, Thermo Fisher Scientific) and PURE system (PUREfrex 1.0, GeneFrontier)^14^. Both cell-free systems use T7 RNAP for transcription. Fluorescence was measured at least for 12 h as a proxy for the expression of sfGFP. All the experiments were performed at least in duplicate.

Figure 1B and C show time-course of GFP fluorescence using several T7 promoter variants including the consensus T7 promoter (denoted as "WT") in the PURE and extract-based system, respectively (See Figure S1 for more details). In the PURE system, the reaction generally took six to nine hours to saturate, while the extract-based system lasted only for three to four hours. A possible reason for this difference can be the rate of DNA and RNA degradation. The PURE system is free from DNase and RNase^31^ (apart from contamination) and therefore the reaction last longer. On the other hand, various nucleases from *E. coli* crude extracts were contained in the extract-based system, which degrade DNA and mRNA and may have caused early termination of protein synthesis.

It is noteworthy that the final expression levels of GFP with many of the variants were higher than that of the consensus promoter in both cell-free systems, while improved protein expression was rather rare in the previous studies with T7 promoter variants in linear form^32^ or plasmid form^26^. To quantitatively compare expression levels between the variants, we normalized the final fluorescence relative to that of consensus promoter. When ranked in descending order (Figure 1D and E), it showed a continuum of protein expression levels in both cases, which is indeed an ideal trait for the fine-tuning of expression level control. Regarding the improved protein expression, 12 and 9 out of 51 variants showed higher expression in the PURE system and extract based system, respectively. The protein expression was increased up to 3.65-fold in the extract-based system. In the PURE system, the maximum increase was less than 1.5-fold.

Although the nearly 4-fold increase in the protein expression seems remarkable, the absolute expression level in the extract-based system was generally quite low compared to that in the PURE system (Figure 4A). We speculate that this low yield would also be due to the nuclease contamination in *E. coli* crude extract as described above. As the core sequence has only five additional base-pair in the upstream of the T7 promoter sequence, the linear DNA templates may have been digested by nucleases and lost the promoter sequence.

However, the extract-based system was excellent at the reproducibility of experiments. Relative protein expression levels of extract-based system were highly reproducible between parallel experiments that were performed on different dates (R^2^=0.97). Those of the PURE system showed a relatively lower reproducibility (R^2^=0.74) (Figure 1F and G). This would be because proteins produced in the PURE system were partially translated or functionally inactive due to ribosome stalling on mRNA^29^. The extract-based systems, on the other hand, contain additional elements that can rescue stalled ribosomes, such as alternative ribosome-rescue factor A (ArfA) with release factor 2 (RF-2)^33^ and elongation factor P (EF-P)^34,35^, and may have contributed to the good reproducibility. A previous study^31^ pointed that the PURE system can produce ~4-fold more protein than an *E. coli* extract-based system. Our results showed the average fluorescence level with PURE system was approximately 3-fold compared to that with the extract-based system (Figure 4A). Considering that some proteins were incompletely translated (and thus no fluorescence), the result seems consistent with the previous study.

Figure 2A and B are heat maps of relative protein expressions using the same data as Figure 1D and E. The promoter strength of each variant showed low correlation between the cell-free systems (Figure 2C, R^2^=0.23). This result was not surprising considering that the extract-based cell-free system contains a number of components that are missing in the PURE system. Plus, DNA/RNA digestion by nucleases from *E. coli* extracts affects mRNA production in the extract-based system. However, the heat maps illustrate that the variants with improved protein expression were mainly located in the upper region of the promoter between the position −17 and −10, corresponding to the polymerase binding domain. This region is particular important for transcription because the T7 RNAP is known to first contact the promoter sequence at position −17 to −13^36^. In contrast, substitutions introduced in the positions between −9 and −5 (PURE) and −11 and −4 (extract-based) significantly reduced the protein expression level. This result is consistent with the previous studies that bases at the position −12 to −5 are crucial for T7 RNAP recognition of the sequence^26,37^. Thus, any substitutions introduced here reduce the protein expression as they interfere contacts with the major groove of the promoter. In the downstream region of the promoter, any substitutions were mostly deleterious to protein expression in both cell-free systems, although protein expression was less affected in the PURE system.

**Figure 2.**
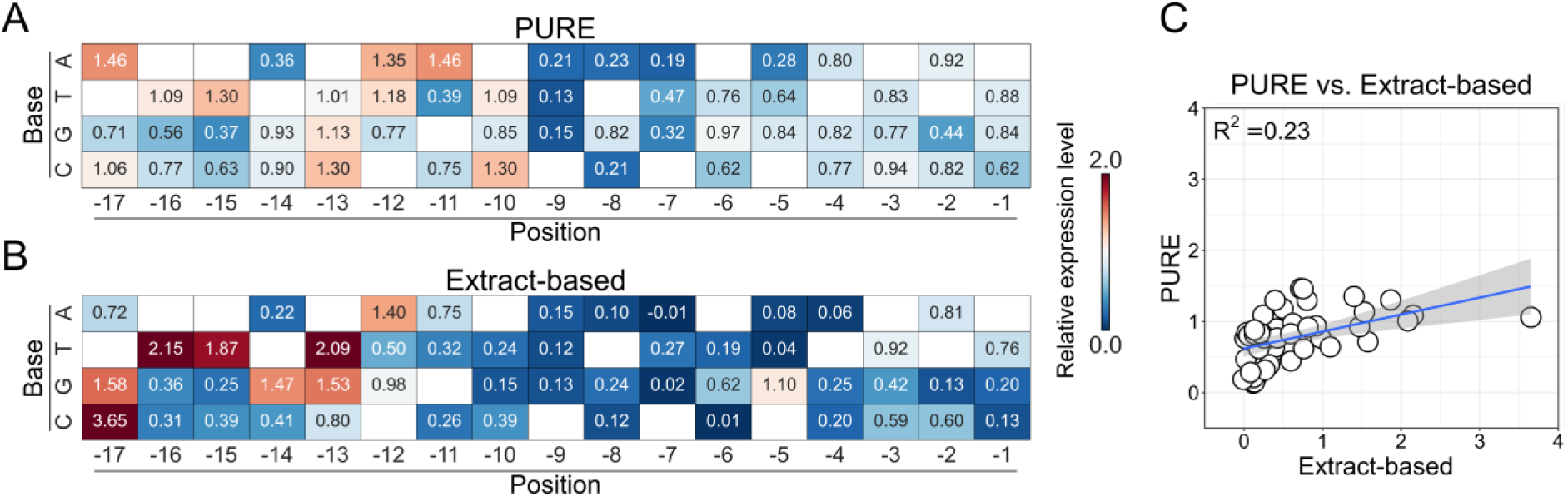
(**A, B**) Heat maps of the relative GFP fluorescence of the 51 T7 promoter variants in the PURE and extract-based systems, respectively. The same data as Figure 1D and E were used. Colors represent fold change in the final expression level relative to that of the consensus promoter. (**C**) Scatter plot of the relative GFP fluorescence for each of the variants. Values obtained with the PURE system were plotted against those with the extract-based system.

### Effect of sequence length on the protein expression

To further investigate the behavior of T7 promoter variants, we constructed another DNA template: Linear DNA with additional sequences attached to the 5’ and 3’-end of the core sequence (hereafter, "extended sequence" for short. Figure 3A). The sequence attached on the 5'-end was a random sequence, whereas that on the 3'-end contained a T7 terminator sequence. We focused on substitutions at position −17 to −10 as they appeared to have diverse effects (beneficial or deleterious) on the protein expression level.

**Figure 3.**
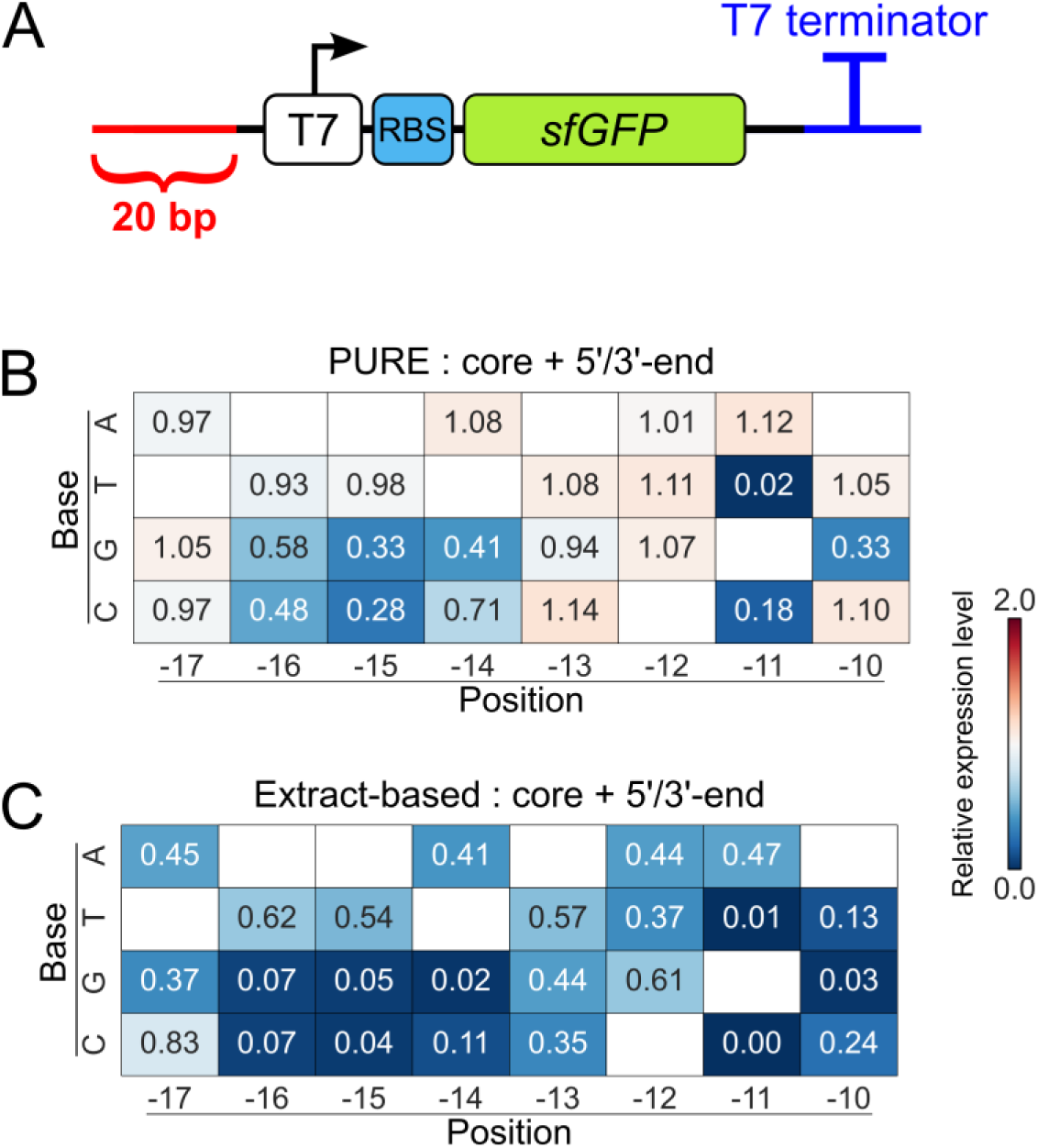
Protein expression in the PURE and extract-based cell-free systems of the T7 promoter variants using a different DNA construct. (**A**) Linear DNA template (core sequence) with extra bases at the 5’-end (red) and 3’-end (blue). The sequence attached at the 3’-end contains a T7 terminator sequence. (**B, C**) Heat maps of the relative GFP fluorescence of the T7 promoter variants with additional sequences attached on 5’ and 3’-ends, evaluated in the PURE and extract-based systems, respectively.

Figure 3B and C show the promoter strength of each variant relative to the consensus sequence. Overall the relative strength decreased in most of the cases compared to the case with the core sequence. In particular, all the variants showed lower protein expression levels than the consensus sequence in the extract-based cell-free system. A possible explanation for the decrease in promoter strength would be more stable binding of T7 RNAP to the linear DNA due to the extra bases: While the substitutions in the −17 to −10 region in the promoter helped the T7 TNAP strap itself more securely to the DNA, the extra bases added at the 5’-end also had the same effect on the T7 RNAP, hence it canceled out the beneficial effect of base substitution. However, the relative promoter strengths of the core and extended sequences showed good correlations in both cell-free systems (Figure S2A and B), suggesting the gene expression levels by the T7 variants are consistent overall.

Although the relative promoter strengths were decreased in the extract-based system, the absolute expression levels by the extended sequence significantly increased more than four-fold on average (Figure 4A and Table S1). In contrast, the expression levels by the extended sequence in the PURE system were similar to those by the core sequence (Table S2). We speculate there would be two possible reasons for the large difference between the two cell-free system: First, additional bases on each end of the extended sequence may have helped prevent the degradation of T7 promoter sequence as well as GFP coding regions by DNase in the extract-based system. On the other hand, the PURE system does not contain nucleases (apart from contaminated ones). Thus, the difference between total protein expressions by the core and extended sequences can be attributed to the dynamics of promoter recognition, which was considered to be relatively minor. Another possible reason for the increase of protein yield in the extract-based system would be due to the T7 terminator introduced on the 3'-end of extended sequence. It has been reported that the T7 terminator improved the stability of transcribed mRNA in *E. coli*-based cell-free system and improved the expression yield more than three-fold^38^. To determine a primary factor of the increase, we constructed core sequence templates with the 5’-end extension. The result showed that the protein expression using the new template was at a similar level as the extended sequence (Figure 5A), suggesting the extended random sequence at the 5’-end was more crucial for the improvement of the maximum expression level. In the case of PURE system, the effect of T7 terminator appeared to be minor (Figure 5B).

**Figure 4.**
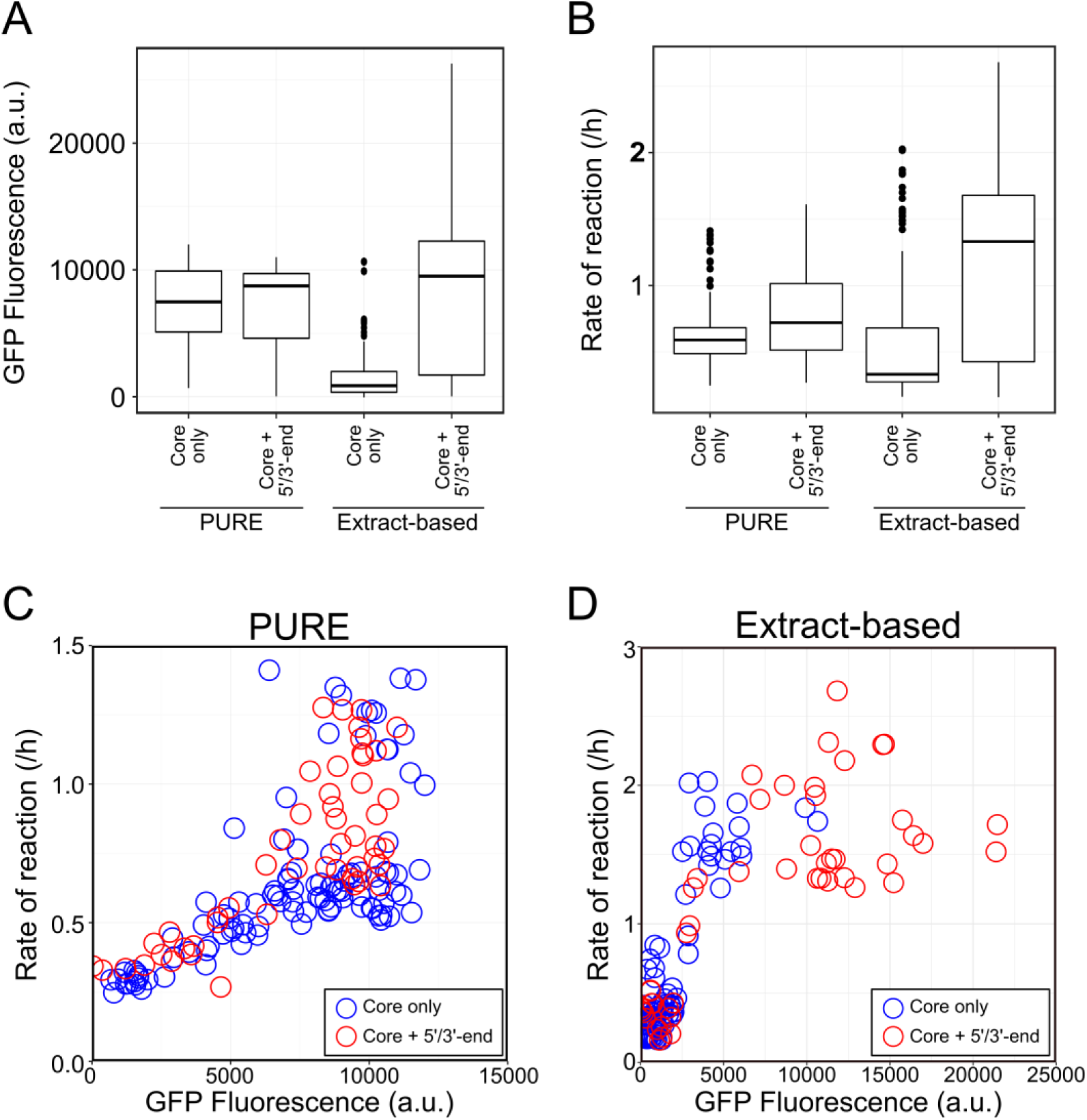
(**A**) Boxplot of absolute GFP fluorescence for different DNA constructs in the PURE and extract-based systems. The black thick line represents the median, and the box shows the first and third quartile. The upper and lower whiskers indicate 50% of the values higher or lower than the median, respectively. Black dots are outliers. (**B**) Boxplot of the rate of reaction from all DNA constructs. The rate was calculated as the slope of a logistic curve fitted to individual fluorescence time-course data. (**C, D**) Scatter plots of the rate of protein expression against the absolute GFP fluorescence for the PURE and extract-based systems, respectively. The same data as (**A**) and (**B**) were used for the plots. Blue and red circles indicate the data for T7 promoter variants with the core and extended sequences, respectively.

**Figure 5.**
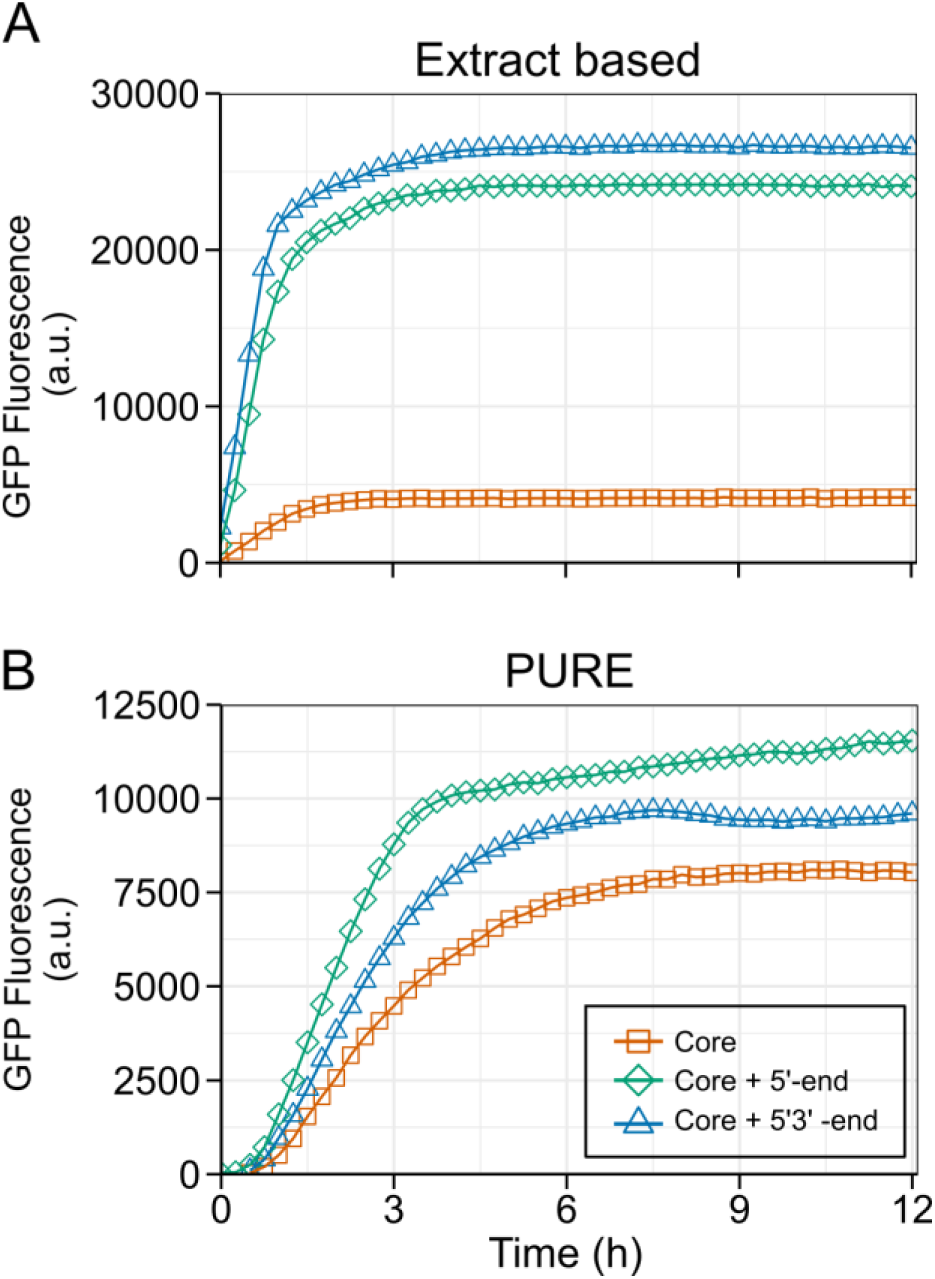
Time-course of GFP fluorescence over a period of 12 h using the T7 consensus sequence. Three different linear constructions, the core sequence (red squares), core sequence with extra bases at 5’-end (green diamonds) and core with extra bases at 5’ and 3’-ends (blue triangles). (**A**) GFP expression in the extract-based system, and (**B**) the PURE system.

Figure 4B shows the rate of protein expression by the core and extended sequences in both cell-free systems. Similarly, to the absolute fluorescence level (Figure 4A), the extended sequence expressed in the extract-based system showed a significant increase compared to the core sequence, whereas a slight increase was observed with the PURE system.

When the data shown in Figure 4A and B were plotted against each other, they illustrated different modes of protein expression dynamics (Figure 4C and D). In the case of PURE system, the scatter plot shows an exponential-like profile: In the low expression region corresponding to protein expressions by weak promoter variants, the absolute fluorescence and the expression rate showed a clear linear correlation. In the high expression region, the promoter variants showed a wide range of expression rates, although it appeared to have a limit of maximum expression level around at 12500. In contrast, the scatter plot for the extract-based system showed a diagonally-rotated profile. The rate of reaction seems to have a limit around at 2 /h while there were a number of variants showing no fluorescence and thus very low rates of expression.

### Modeling two cell-free systems

We found the results in Figure 4C and D particularly interesting because the system parameter we varied here was the binding affinity of T7 RNAP to a promoter sequence (corresponding to the transcription constant *k*_*TX*_), but the responses of two cell-free systems to the change were very different.

To investigate possible dynamics behind the behaviors pf two cell-free systems, we fitted each GFP measurement time-course curves with the equations (1)–(3) shown above and estimated the parameter values, *k*_*TX*_, *k*_*TL*_, *d*_*D*_, *d*_*R*_ and *d*_*p*_. Figure 6 shows the histograms of the fitted parameter values. Note that a modified equation was used to fit the PURE system data because of the reason described below. For fitting of the extract-based system, the above equation was used.

**Figure 6.**
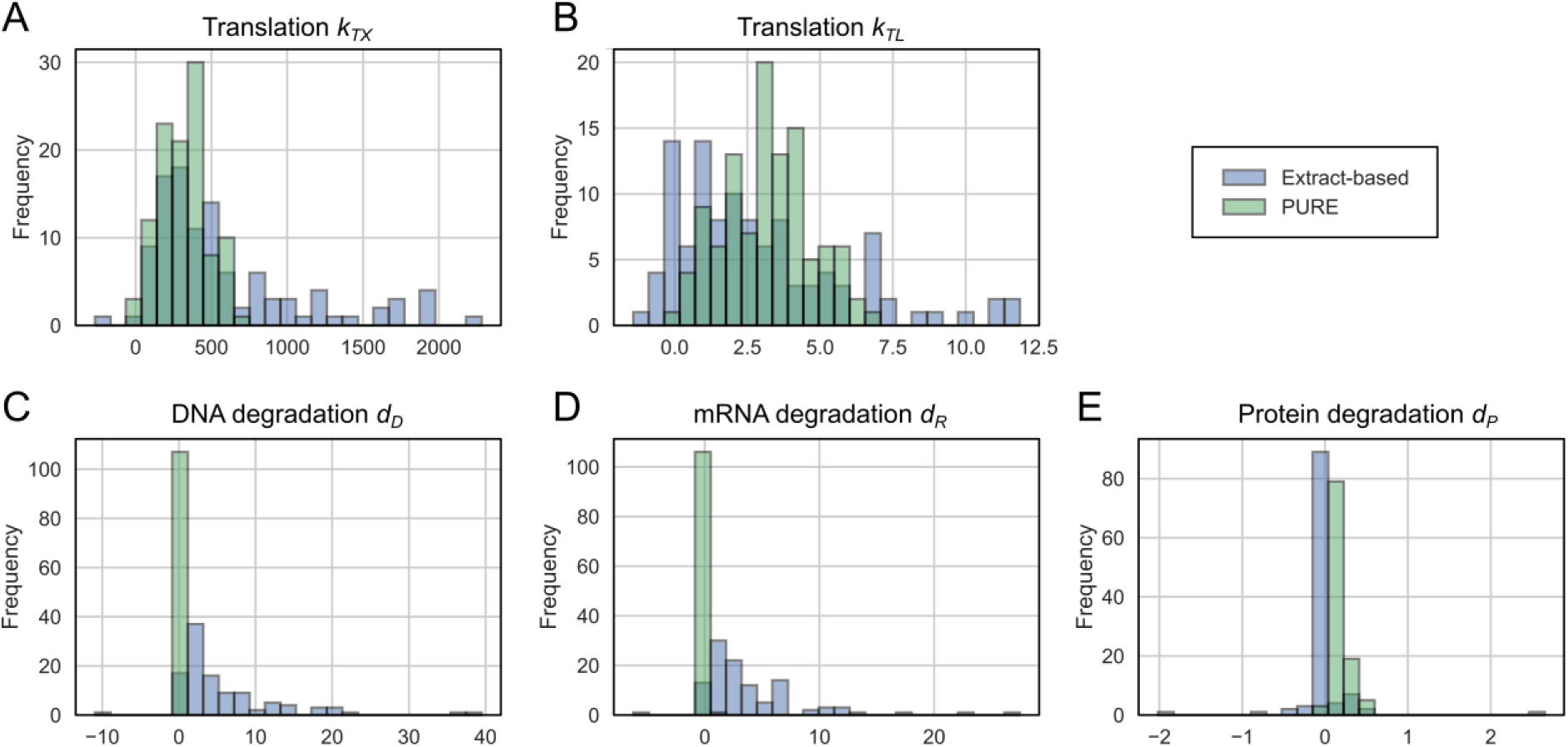
Histograms of estimated parameter values. (**A**) Transcription constant *k*_*TX*_, (**B**) translation constant *k*_*TL*_, (**C-E**) degradation constants for DNA *d*_*D*_, RNA *d*_*R*_, and protein *d*_*p*_, respectively. Note that different models were used to fit the parameters for the PURE and extract-based cell-free systems. See text for details.

The fitted parameter values were indeed consistent with the possible mechanisms behind the observed protein expression as we discussed above. The transcription parameter *k*_*TX*_ showed a wide distribution over three orders of magnitude due to the altered binding affinity by substituted bases in the T7 promoter sequence. In contrast, the distribution of the translation parameter *k*_*TL*_ was much narrower and most of the parameters fit within the same order of magnitude. The DNA/RNA degradation terms, *d*_*D*_ and *d*_*R*_, showed a large difference between the two cell-free systems. While those for the extract-based system showed a wider distribution, those for the PURE system had only one single peak. They indicate that DNA and RNA are very slow to degrade in the PURE system because it does not contain nucleases, whereas they can be quickly digested in the extract-based system. In both systems, protein degradation term *d_p_* was close to zero. This is consistent with the experimental data as the measured GFP fluorescent signal did not decrease over a long period of time (see Figure S1).

Next, we simulated the cell-free protein expression using the fitted parameter values and equations above to study the internal dynamics of cell-free protein expression. From the systems point of view, altering the promoter sequence corresponds to varying *k*_*TX*_ because substituted bases affect the binding affinity of T7 polymerase to the promoter. To simulate the protein expressions by the promoter variants, we used the parameter values fitted for the consensus promoter sequence and varied only *k*_*TX*_ from zero to the maximum fitted value.

Figure 7A shows simulated protein expression in the extract-based system. It well reproduces the protein expression in the cell-free system, such as halting of protein expression after 3-4 h. When calculated the rate of reaction and maximum protein expression level by fitting the simulated data to a logistic curve, it also showed a similar profile as shown in Figure 4D.

**Figure 7.**
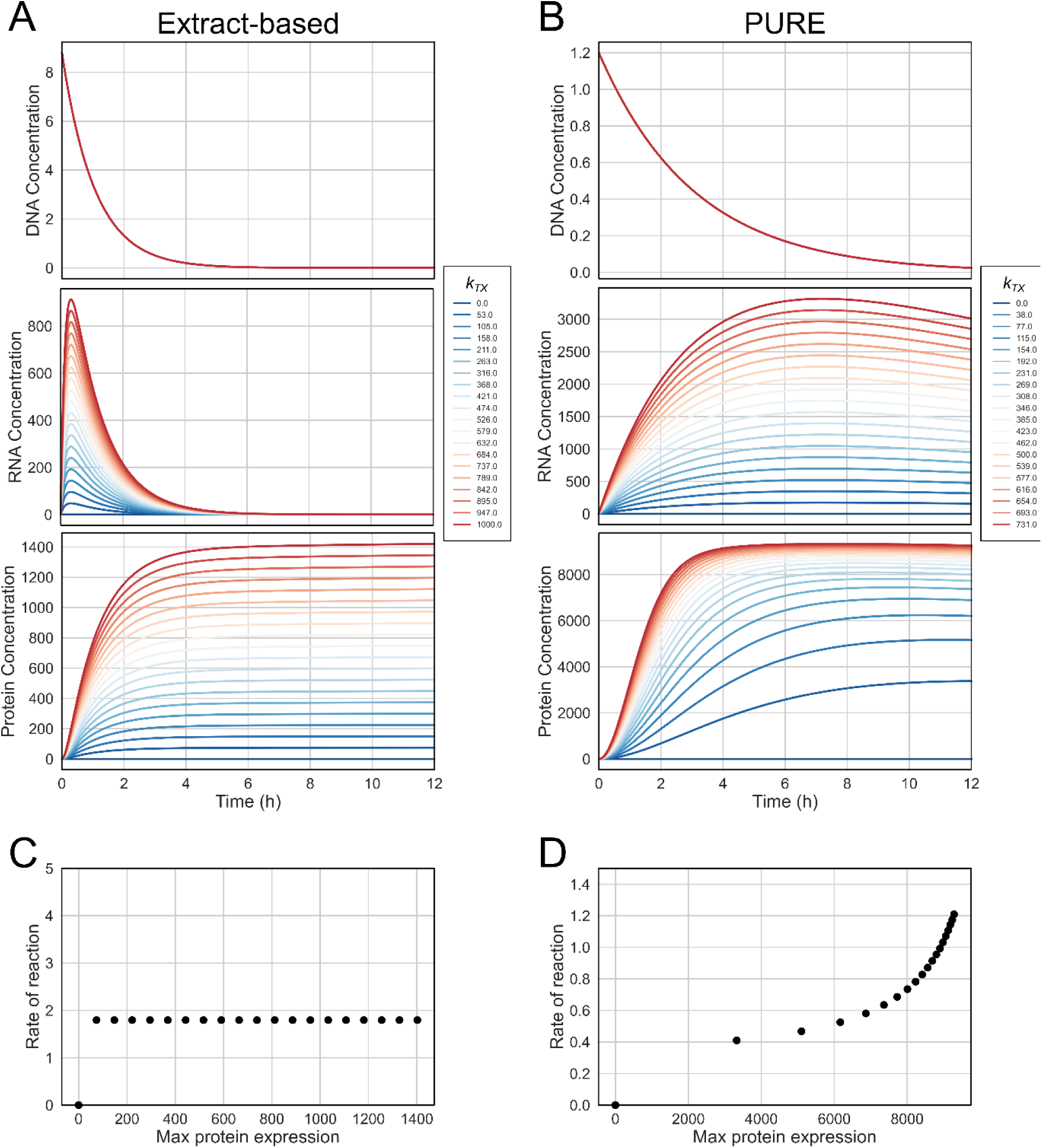
Simulated cell-free protein expression. (**A**) Simulated cell-free protein expression of the extract-based system. DNA concentration (top), mRNA concentration (middle), and protein expression (bottom) were shown. The parameter *k*_*TX*_ was varied (colors) while the other parameters were fixed. Simulated cell-free protein expression of PURE system. A modified equation (see text for details) was used to simulate protein expression. (**C, D**) Scatter plots of the rate of protein expression against the maximum protein expression in the extract-based and PURE systems, respectively. The values were obtained by fitting the simulated data in (**A, B**) to a logistic curve.

We performed the same simulation using the fitted parameter values for the PURE system (Figure S3). Although the temporal expression patterns simulated the observed temporal expression profiles, it did not reproduce the unique scatter pattern shown in Figure 4C. This result implies that there were some factor(s) missing in the above theoretical model.

It is widely recognized that protein synthesis in a cell-free system is limited by the availability of energy source and accumulation of inhibitory byproducts^31^. In fact, the protein expression yield significantly increases up to 72-fold by feeding energy molecules and removing byproducts through dialysis membranes^39^. It was also shown that efficient recycling of inorganic phosphate improves the total protein yield in a cell-free system^13^. From the systems point of view, this can be considered as negative feedback to the protein production. To incorporate this factor, we modified the equation (3) as follows:

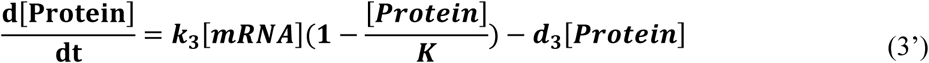

The term (1 − [*Protein*]/*K*) represents the negative feedback. The proteins ceases to produce as the protein expression level reaches the maximum defined by the carrying capacity constant *K*. Using the equation (1), (2), and (3'), we re-fitted the model to experimental data to estimate the system parameters *k*_*TX*_, *k*_*TL*_, *d*_*D*_, *d*_*R*_, *d*_*p*_, and the carrying capacity *K* (see Figure 6). Figure 7B shows the simulation results performed with varying *k*_*TX*_. While the protein expression profile is similar, the exponential-like profile of the rate of reaction against the maximum expression level was also simulated (Figure 7D).

When the simulation results in Figure 7A and B were compared, there are a few clear differences. First, the time-course profiles of RNA concentration are quite different in the two cell-free systems. In the extract-based system, mRNA was produced very rapidly, which peaked around at 30 min after the start of the reaction. The concentration of mRNA decreased exponentially after the peak as the template DNA was degraded by nuclease present in the reaction mixture, and eventually reached to the base level around at 4 h. In the PURE system, the template DNA also degraded but the rate was much slower than the extract-based system. As a result, the rate of mRNA production was slower than the extract-based system, but the most of transcribed mRNA were present even after 12 h.

We experimentally confirmed the temporal dynamics of RNA in both cell-free systems (Figure 8). To do this, we constructed a short DNA template consisting of a T7 promoter, RBS, and a Spinach aptamer. The Spinach RNA aptamer shows green fluorescence in the presence of fluorophore, such as 3,5-difluoro-4-hydroxybenzylidene imidazolinone (DFHBI)^40,41^. Because of this green fluorescence, we did not include sfGFP gene in the DNA template as the peak emission wavelength of Spinach aptamer was quite close to that of sfGFP. We tested with the consensus T7 promoter sequence, two strong variants, as well as a weak variant. The temporal profiles of the measured mRNA concentration were similar to the simulated mRNA concentration in both cell-free systems. It should be noted that no experimental information about mRNA concentration was included when fitting the differential equations to the protein expression data. Nevertheless, the time-courses of mRNA concentration in the simulation reproduced those in the experiments. This result, in turn, validates that the theoretical models capture the dynamics behind the cell-free protein expression.

**Figure 8.**
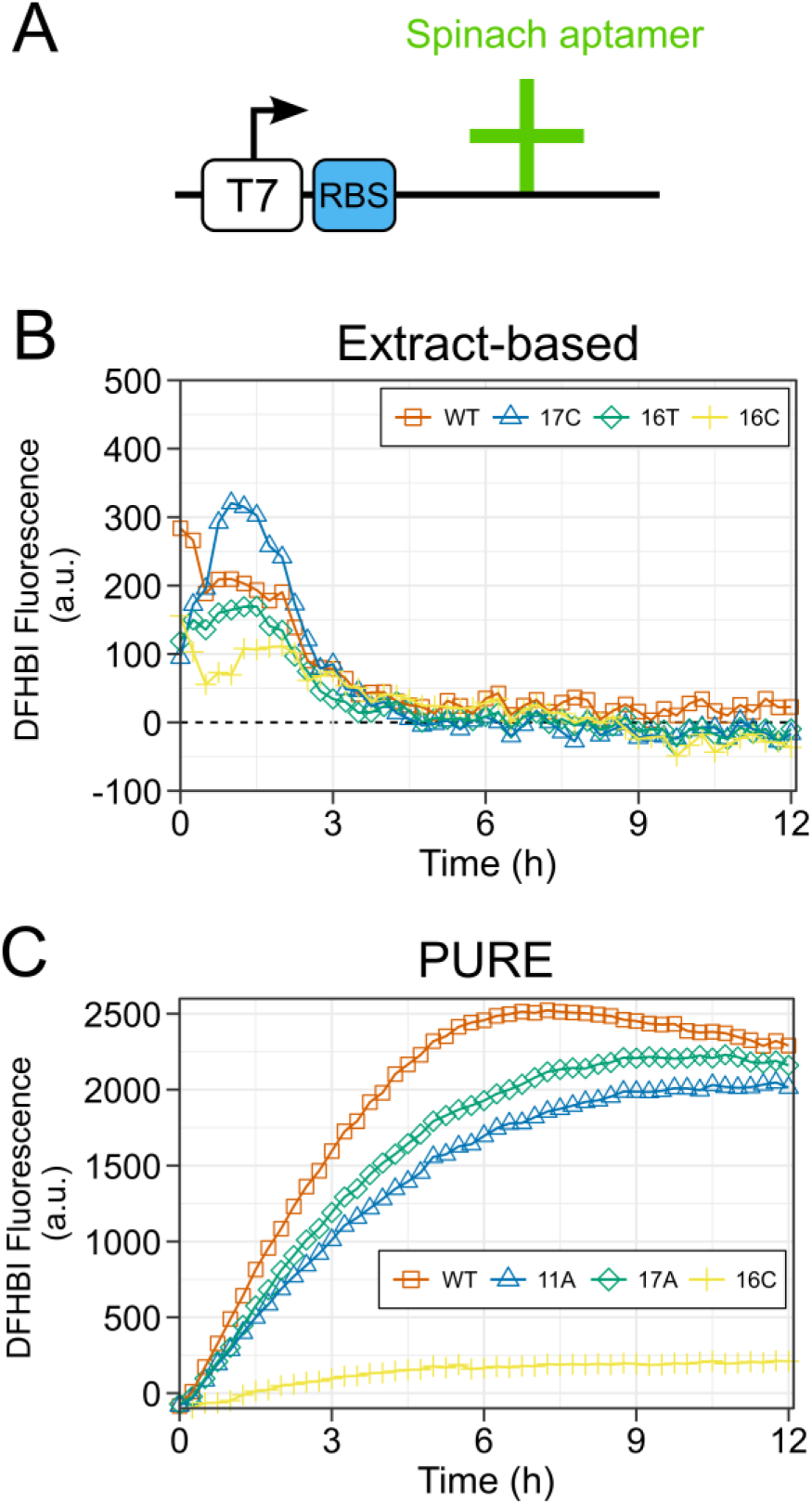
(**A**) Linear DNA template with Spinach aptamer. The sequence contains the T7 promoter or its variant, RBS, and Spinach aptamer. It does not contain any genes to be expressed. (**B, C**) Fluorescence measurements of Spinach aptamer with the consensus promoter sequence (denoted as WT) and other three variants in the PURE and extract-based systems, respectively.

Another difference between the two simulations was the response of cell-free system to a change in *k*_*TX*_ (Figure 7A and B bottom figures for protein expression). When the parameter *k*_*TX*_ linearly varied, the protein expression yield in the extract-based system also linearly changed because there were no nonlinear terms in the model equation. In the PURE system, the response of the system to the linear change was nonlinear because of the negative feedback term. As a result, the system was very sensitive to changes in the smaller range of parameter values, which implies that the maximum expression levels tend to be biased to the higher region. Indeed, histograms of the experimental and simulated maximal expression levels (Figure S4) showed similar distributions in the case of PURE system, i.e. the distributions were biased to higher expression range. Combined time-course plots for both cell-free systems (Figure S5) showed similar patterns as the simulated cases in Figure 7, especially the cases with the extended sequences. Although we were not able to experimentally measure *k*_*TX*_ (the binding affinity of T7 RNAP to each promoter variant sequence), these results may also validate the models.

These observations illustrate that the two cell-free systems have different limiting factors for improved protein expression. In the extract-based system, the protein expression was primarily limited by the availability of template DNA and transcribed mRNA. As they were quickly digested by nucleases in the reaction mixture, any ways to prevent or slow down the degradation would improve the yield. This is consistent with the experimental results above that the extended sequence showed significantly improved yield. In the PURE system, the limiting factors lie within the mRNA-protein translation. In the current form, the negative feedback term in the model is still ambiguous and needs further biological elaboration. However, it has been shown that supplementing amino acids, tRNAs, magnesium, and proteins involved in ribosome recycling to a PURE system improves the yield, which is consistent with our model prediction.

In this work, we characterized the behavior of single base-pair substituted variants of the T7 promoter in two different cell-free systems. Although further understanding of cell-free protein expression mechanism at the molecular level is clearly required to fully explain the complex expression patterns observed here, our experiments revealed different modes of protein expression dynamics in the PURE and extract-based systems, which were confirmed by model analysis. Our result would illustrate that even simple time-course measurement data of cell-free systems contains rich information and the combination with systems approach help us uncover the dynamics behind it. In fact, the systems approach has already proven effective for engineering synthetic genetic circuits in the cell-free system^24,42^. We believe that this approach can also be effective to disentangle the complex dynamically-interacting factors in the cell-free system and obtain deeper insights that are otherwise difficult to capture.

## METHODS

### Materials

All the chemicals and materials were purchased from Thermo Fisher, except for the following: the Zyppy Plasmid Miniprep Kit, ZymoPURE Plasmid Midiprep Kit, and Mix and Go *E. coli* and Transformation Kit and Buffer Set were acquired from Zymo Research; the Wizard SV Gel and PCR Clean-Up System and Wizard *Plus* SV Minipreps DNA Purification System were bought from PROMEGA; (5Z)-5-[(3,5-Difluoro-4-hydroxyphenyl)methylene]-3,5-dihydro-2,3-dimethyl-4H-Imidazol-4-one (DFHBI) fluorophore was bought from Sigma Aldrich. Cell-free PURE system (PUREfrex 1.0) was bought from GeneFrontier Corporation and the Expressway system (Expressway™ Maxi Cell-Free E. coli Expression System) was bought from Thermo Fisher Scientific. All the primers were custom-ordered from Thermo Fisher Scientific. All the PCR reactions were performed using KOD Hot Start Master Mix from EMD Millipore. All the PCR products were verified for size in a 1.5% agarose gel, then cleaned using the Wizard SV Gel and PCR Clean-Up System and subsequently measured with Nanodrop for DNA quantification.

### Linear DNA constructs

The plasmid sfGFP-pET32b, containing sfGFP as the reporter gene, was used as the template to create the genetic constructs. *E. coli* BL21-CodonPlus(DE3)-RIPL competent cells were transformed with the plasmid sfGFP-pET32b following the protocol specified for the competent cells, and the colonies selected in LB agar plates with 50 µg/ml of ampicillin. The LB agar plates were grown overnight at 37°C before selecting the colonies. The chosen colonies were incubated in 10 ml of 2xYT medium with 50 µg/ml of ampicillin, overnight at 37°C and 200 rpm. The plasmid sfGFP-pET32b was extracted from the bacterial culture using the Zyppy Plasmid Miniprep Kit. The plasmid was then used to make a linear construct of the sfGFP gene with a RBS region and T7 promoter.

Linear DNA constructs containing variants of T7 promoter with RBS and sfGFP were constructed with two-step PCR by following the protocol for PUREfrex 1.0: The first PCR performed was on the plasmid, sfGFP-pET32b, to add the RBS region using the primers RBS-sfGFP-F and sfGFP-R (Supplementary Table 1). The PCR conditions are as follows: Initial Denaturation (94°C, 120 sec), 35 cycles of denaturation (98°C, 10 sec) and annealing/extension (68°C, 30 sec), and Final Extension (68°C, 120 sec). The results construct, RBS-sfGFP, was then used for a second round of PCR using the primers T7-RBS-sfGFP-F and sfGFP-R (Supplementary Table 1) with the following conditions: Initial denaturation (94°C, 120 sec), 35 cycles of denaturation (98°C, 10 sec), annealing (30°C, 30 sec) and extension (70°C, 30 sec), and final extension (70°C, 120 sec). The result construct, T7-RBS-sfGFP containing the consensus T7 promoter sequence, was used throughout the experiments as reference. It was also used as the template DNA to construct the different T7 promoter variants. A different forward primer was used for each variant (Supplementary Table 1) but the reverse primer, sfGFP-R, was the common for all of them. For making all the variants, the PCR conditions were: Initial denaturation (95°C, 120 sec), 35 cycles of denaturation (95°C, 20 sec), annealing (67°C, 10 sec) and extension (70°C, 15 sec), and final extension (70°C, 120 sec).

Using the T7 variant linear DNA constructs, extra bases were added only at 5’ end and both at 5’ and 3’ ends. The sequence attached at the 5’ end was non-coding random sequence, and that at the 3’ end contains a T7 terminator sequence. For the samples with added bases at 5’ and 3’, specific forward primers for each variant were used along with a common reverse primer, 5p3p_Common_R (Supplementary Table 2). The PCR conditions were: Initial denaturation (95°C, 120 sec), 40 cycles of denaturation (95°C, 20 sec), annealing (56°C, 10 sec) and extension (70°C, 30 sec), and final extension (70°C, 10 min). For the WT sample with extra bases only at 5’ end, its specific forward primer was used as described above with the difference of the reverse primer, 5p_Common2_R (Supplementary Table 2). The PCR conditions were the same as described before.

### Spinach RNA aptamer constructs

Linear DNA templates containing the spinach aptamer was constructed using four primers: Aptamer1-F, Aptamer2-F, Aptamer3-R and Aptamer4-R (Supplementary Table 2). The PCR conditions were: Initial denaturation (95°C, 120 sec), 40 cycles of denaturation (95°C, 20 sec), annealing phase (62°C, 10 sec) and extension (70°C, 5 sec), and final extension (70°C, 10 min).

The PCR product was purified and then used as a template for another PCR. A forward primer that was used to make a core sequence construct (Supplementary Table 1) and a common reverse primer Aptamer4-R were used. Following PCR conditions were used: Initial denaturation (95°C, 120 sec), 40 cycles of denaturation (95°C, 20 sec), annealing phase (57°C, 10 sec) and extension (70°C, 5 sec), and final extension (70°C, 10 min).

### Cell-Free transcription-translation reaction

Unless otherwise indicated, all the reactions using the PURE system were performed with a final volume of 20 µl and using 24 ng (1 µl) of each linear DNA construct. For the reactions using the Expressway system, the total volume of each reaction was 26.6 µl with 235 ng (5 µl) of linear DNA. Stocks of DNA samples were made for reproducible experiments and all experiments were performed in duplicate. The reaction components for both systems were assembled in a master mix for each one and then added to the corresponding amount of DNA in a black flat-bottom 364 well-plate (Nunc 384 black well-plate, Fisher Scientific). The well-plates were covered with a transparent sealing membrane (Breath-Easy) to avoid evaporation and then incubated in a plate reader (Infinite 200 PRO, Tecan) at 37°C for 12-20 h. During the incubation period, GFP fluorescence was measured and recorded every 15 min (excitation: 395 nm; emission: 509 nm).

For the RNA spinach aptamer experiments, DFHBI fluorophore was added to each reaction to a final concentration of 20 µM and the DFHBI fluorescence measured (excitation: 460 nm; emission: 502 nm).

### Mathematical analysis and modelling

Computer simulation was performed using custom programs written in Python and R^43^. For numerical simulation of differential equations, SciPy module^44^ was used. Nonlinear fitting of experimental data for the estimation of system parameters in the differential equations were performed by least square fitting using leastsq function in the SciPy module. The rate of reaction and the maximum expression level for each time-course fluorescence data were calculated in R using growthcurver package^45^.

## Associated Content

### Supporting Information

The Supporting Information is available free of charge on the ACS Publications website at DOI: ??????.

> All the time-course plots of GFP expression from T7 promoter variants, correlations between the core and extended sequences, simulated cell-free protein expression in the PURE system using the standard model, histograms and kernel density plots of maximum protein expression levels comparing experimental and simulated data, time-course of GFP fluorescence, fold change of the maximum GFP fluorescence, and a list of primers used in this study.

## Notes

The authors declare no competing financial interest.

## Acknowledgements

This work was supported by the Lord Kelvin Adam Smith fellowship and PhD scholarship from The University of Glasgow, JSPS KAKENHI (16H06156 and 16H00797) and Astrobiology Center Project of the National Institutes of Natural Sciences (NINS) (AB291017). This project was made possible in part through the ELSI Origins Network (EON)'s Seed Grant, which is supported through a grant from the John Templeton Foundation. Authors thank Dr Kazufumi Hosoda for their comments on the early version of the manuscript.

